# Ribosomal DNA revealed an extensive role of allopolyploidy in the radiation of *Ulex* L

**DOI:** 10.1101/2021.01.20.427424

**Authors:** João P. Fonseca, Ana Pereira, Joana I. Robalo, Carlos Neto, José C. Costa

## Abstract

We studied the phylogeny of *Ulex* L., a genus of spiny legumes, which its center of diversity in the Iberian Peninsula, using ribosomal DNA markers (rDNA), namely ETS, 5.8S and ITS (45S), and 5S intergenic spacer regions. One of the main findings was the presence of very different haplotypes in 5S-IGS genes and, to a less extent, in ETS and ITS, in seven polyploid *taxa.* We interpreted these results as an indication of hybrid origins and proposed allopolyploidy for *U. argenteus* ssp. *subsericeus, U. australis ssp. australis, U. australis* ssp*. welwitschianus, U. densus, U. europaeus* ssp. *europaeus, U. europaeus* ssp. *latebracteatus,* and *U. jussiaei.* These results reinforce an early hypothesis which stated that the radiation of *Ulex* occurred mainly by polyplodization.

Phylograms showed two main clades, one grouping the hydrophilic *U. gallii*, *U. breoganii* and *U. minor*, and the other grouping the southern, xerophytic, *taxa*. The putative allopolyploids showed haplotypes, which grouped in both clades, indicating that allopolyploidy, occurred through hybridization from these hydrophilic and xerophytic lineages.

The phylogenetic position of *U. micranthus* is not certain and it is discussed. The 5S-IGS showed to retain more polymorphisms than ETS gene or ITS markers. This result is compatible with the hypothesis that 5S rDNA region is less vulnerable to inter-loci concerted evolution than 45S, providing a more suitable marker for reconstructing histories of allopolyploid species. We discuss the taxonomic consequences of these results.

## Introdution

Polyploidy is widely acknowledged as a major mechanism of speciation in plants. It can lead to speciation in several ways, either within single individuals or following hybridization between closely related populations (autopolyploidy), or from interspecific hybridization events (allopolyploidy) (1).

Nuclear ribosomal DNA has proved to be a valuable instrument to investigate polyploidy events. In eukaryotes, nuclear ribosomal DNA (rnDNA) belongs to two universal gene families: the 5S nrDNA and 45S nrDNA. Both are organized in tandem arrays. In most plants, they have different locations in the genome (2,3). The 45S array is widely utilized in phylogenetic studies, and is composed by an extensive intergenic spacer (IGS), an external transcribed spacer (ETS), three genes encoding the ribosomal units for 18S, 5.8S, and 28S, and two internal transcribed spacers (ITS1 and ITS2) (4). The other tandem repeat codes 5S nrRNA, and includes a small intergenic spacer (5).

After allopolyploidization, ribosomal DNA can undergo two processes, which can lead to the loss of parental haplotypes: concerted evolution and the loss of loci (6,7). Concerted evolution is a process which homogenizes the sequences within a species, making them more similar to each other than to the homologous sequences of other species reducing haplotypic variability. Some authors reported differences between the concerted evolution for the 45S and 5S arrays. Evidence was presented suggesting that, in 45S, the homogenization mechanisms operate within loci as well as between different rnDNA loci, while in 5S concerted evolution only operates within loci or, at least, homogenization mechanisms inter-locus are not very effective (3,8,9).

If concerted evolution is not completed, a hybrid species can retain two or more very divergent haplotypes originating from each parental *taxa* (10). If these haplotypes are found in other lineages in homozygosity this is often considered a consistent clue of a hybrid origin, and it is used to identify parental lineages (11,12). However, the presence of a single haplotype in a polyploid do not mandatorily indicates autopolyploidy, because this can also arise when the concerted evolution is complete and one of the haplotypes of the parental species is lost (13).

The *Ulex* L. genus belongs to the *Fabaceae* family and *Genisteae* Benth. tribe. It is native of Western Europe and northwest Africa and Iberian Peninsula has been considered as its center of diversity (14). They are spiny shrubs, characterized by the presence of leaves reduced to small spines (15). This genus includes more than 20 *taxa* (16,17), considering species and subspecies, and polyploidy is frequent. Following Cubas (18) and Cubas et al. (15), the genus includes, at least, 12 polyploid taxa, including tetraploids and hexaploids (Table 1).

**Table 1.**
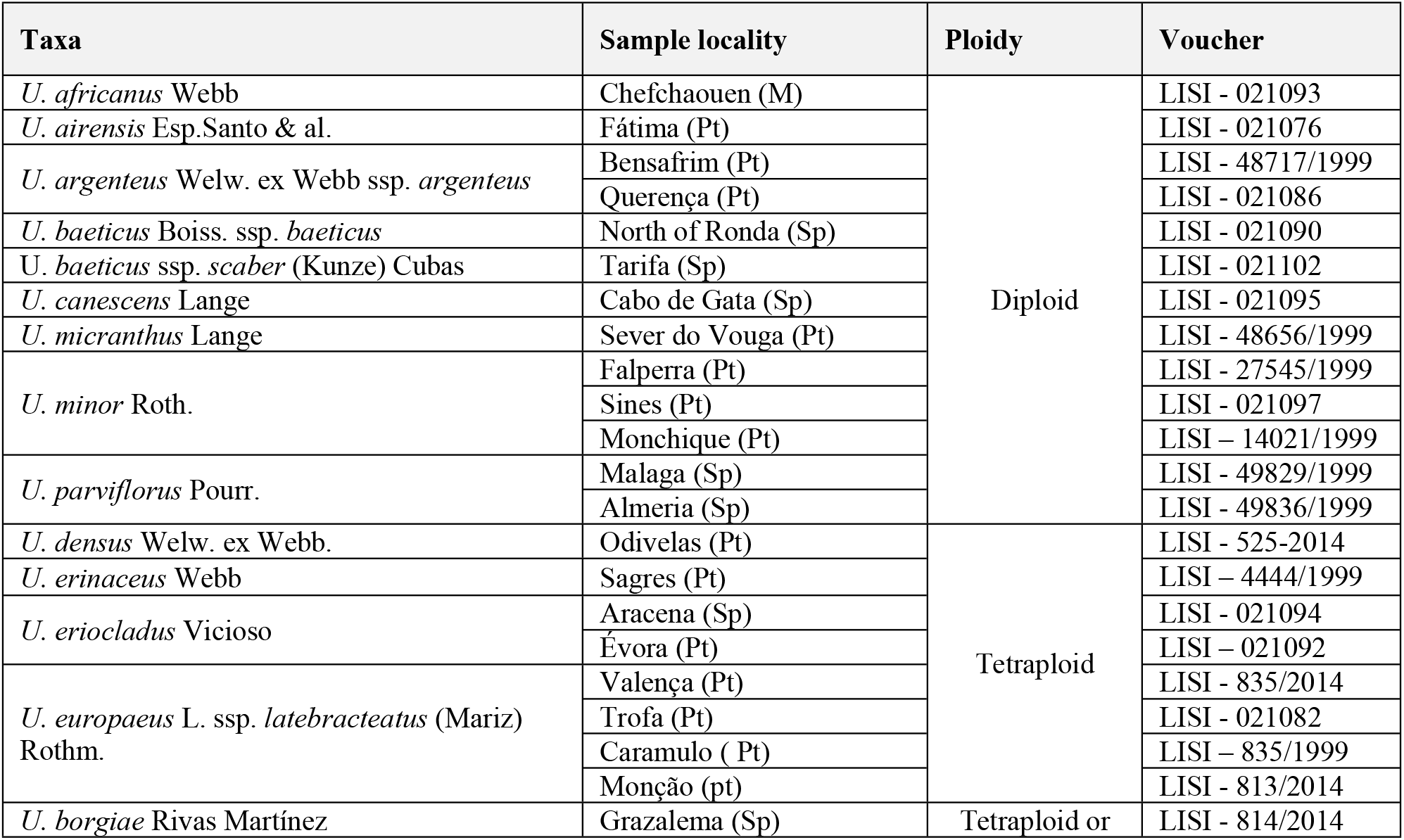
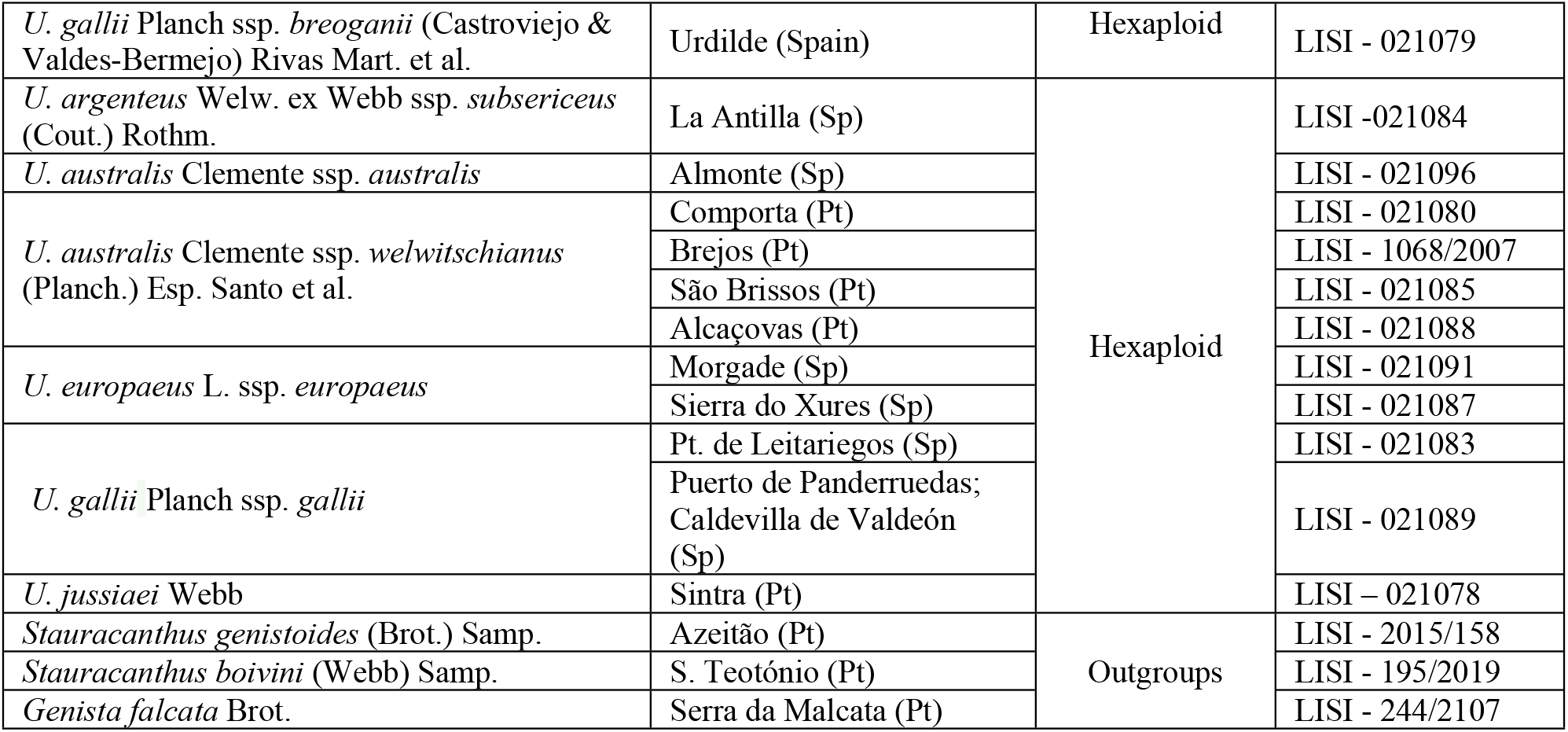
Location, ploidy, and voucher of the studied plants. Sp – Spain; M – Morocco; Pt – Portugal. Vouchers were deposited in the Herbarium of Instituto Superior de Agronomia (Lisbon).

Four previous studies tried to clarify the phylogenetic relationships of this genus, using the internal transcribed spacers of nuclear ribosomal DNA (nrDNA) and plastid markers. Ainouche et al. (14) found that *Ulex* is monophyletic and indicated that *U. micranthus* Lange is a monospecific lineage, separated from all other species. Similar results were achieved by Pardo et al. (19) with the same markers. However, all these markers showed low variability and generated poorly resolved phylograms (19). Ainouche et al. (14) suggested that the low level of sequence divergence reveals a recent radiation and that polyploidization could have played an important role in the speciation process (14). Cubas et al. (15), using chloroplast microsatellite markers, proposed an evolutionary scenario for the evolution of Moroccan and Andalusian (Spanish) species. According to these authors, Southern Iberian diploid *taxa* were closely related and the polyploid *U. borgiae* Rivas Martínez arose from the hybridization of *U. baeticus* Boiss. ssp. *baeticus* and *U. baeticus* ssp. *scaber* (Kunze) Cubas. Finally, Dias et al. (20) presented sequences of nrDNA of *Ulex* and advocate that *U. europaeus* L. subsp. *europaeus* is an allopolyploid (20). The knowledge on the phylogenetic relationships does not go much further.

The main goal of this paper is to study the phylogeny of *Ulex.* Considering the high number of polyploid *taxa* and the low genetic variability reported, we tested the hypothesis of Ainouche et al. (14), which stated that the radiation within this genus resulted mostly from polyploidy events, including allopolyploidy. We used three nuclear fragments of nuclear nrDNA, two from the 45S arrays, namely external transcribed spacer (ETS) and a fragment with two internal transcribed spacers (ITS1 and ITS2), the gene 5.8S (henceforward called ITS), and the 5S intergenic spacer (5S-IGS).

## Material and methods

### Plant material

Our sampling covered most of *Ulex* taxa, at least those classified as independent species, except the Moroccan *U. congestus.* Thirty-three populations from twenty-one taxa were examined (Table 1). The sequences of *U. europaeus* ssp. *europaeus* of 45S rnDNA (ETS and ITS) were retrieved from GenBank (Accession numbers: KC806125.1 to KC806140.1, and KC832359.1 to KC832370.1). *Genista falcata* Brot., *Stauracanthus genistoides* (Brot.) Samp. and *Stauracanthus boivinii* (Webb) Samp., were used as outgroups. Sequences were deposited in GenBank (available at www.ncbi.nlm.nih.gov/). Samples consisted of fragments of phyllodes. Vouchers were deposited in the Herbarium of Instituto Superior de Agronomia (Lisbon). We also used preserved material from this herbarium. Criteria for morphological identification followed Cubas (18).

### DNA extraction, amplification, sequencing and editing

DNA was isolated using the MasterPure Plant Leaf DNA Purification Kit (Sigma-Aldrich Corporation). The PCR mix was prepared for a volume of 20 μl, with 10 μl of RedExtract-N-ampl PCR reaction mix (Sigma-Aldrich), 0.8 μl of each primer (10 M), 4.4 μl of sigma-water and 4 μl of template DNA.

In a first scoping, the marker 5S-IGS was amplified with the primers S1 and AS1 (21). At this step, some polyploids presented chromatograms with double peaks or long illegible fragments. In these cases, we design allele-specific primers (Table 2), a technique that uses primers with deliberate mismatches towards one of the template copies to make the reaction specific for one of the templates (22,23). It was impossible to design allele-specific primers for the 45S fragments (ETS and ITS), as well as to split two haplotypes of 5S-IGS of *U. argenteus* Welw. ex Webb ssp. *subsericeus* (Cout.) Rothm., mainly due to few variable sites. In these cases, chromatograms with double pics were read by the method described by Sousa-Santos et al. (24), using the indels (24).

**Table 2.**
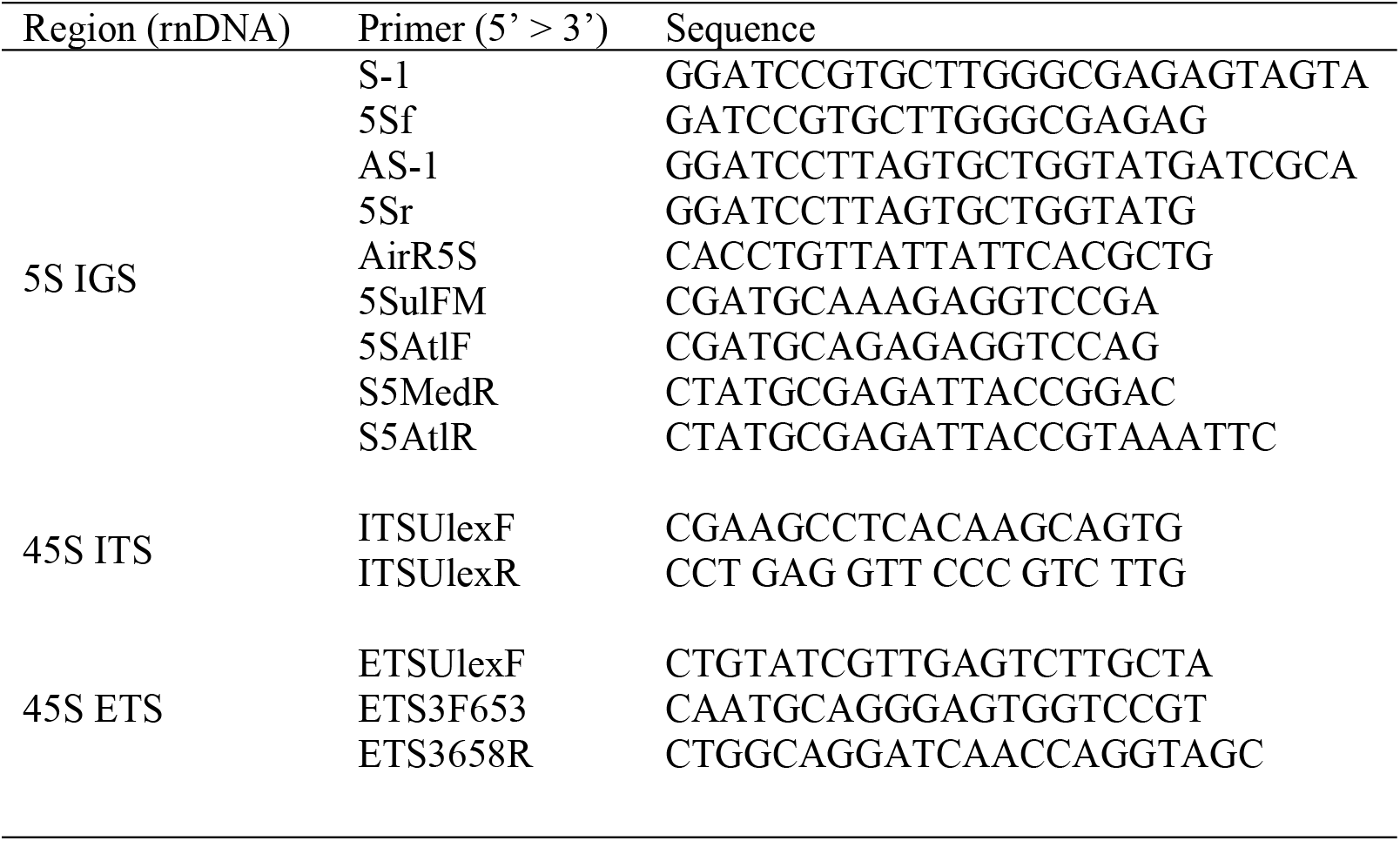
Primers for PCR amplification. S-1 and AS-1 following Sun et al. (21). The other primers were designed for this study

All fragments were amplified with the same PCR protocol. PCR’s included an initial denaturation of 2 min at 94° C, followed by 35 cycles of 1 min. denaturation (94° C), 45 s. annealing (53° C), 50 s. elongation at 72° C, and a final extension at 72° C (7 min.). PCR products were purified with the Microclean kit (Microzone Ltd), and sequenced using the same primers (in STABVIDA, Portugal).

DNA sequences were edited using BioEdit v. 7.0.1 (25) and aligned using ClustalW (1.81) (26), with default gap opening penalties.

### Phylogenetic Analyses

Reconstruction of phylogenetic relationships was conducted using Maximum Parsimony (MP), and Bayesian Inference (BI). The BI was performed in Mr. Bayes 3.1 (27) using MCMC, with two independent runs of four Metropolis-coupled chains of two million generations each, to estimate the posterior probability distribution. Topologies were sampled every 100 generations, and a majority-rule consensus tree was estimated after discarding the first 2000 generations sampled. Sequences were partitioned by region (ITS and ETS), in the concatenated matrix. The MP analysis was conducted in PAUP 4.0b10 (28), using a heuristic search with random stepwise addition (1000 replicates), tree-bisection-reconnection branch swapping. All characters and character-state changes were equally weighted. Bootstrap analysis (1000 replicates) was used to assess the robustness of branches of the MP trees (29). The best-fit model of nucleotide evolution was obtained using Jmodeltest 3.7 (30) with the Akaike Information Criterion (AIC) (31).

Indels shared by two or more haplotypes were coded as binary characters according to the ‘‘simple indel coding” method of Simmons & Ochoterena (32) and used in parsimony and Bayesian searches (32). The binary gap data and the sequence data were treated as separate partitions.

For most taxa it was possible to concatenate the sequences of ETS and ITS, because we found only one haplotype or very similar ones from each marker, for each plant. This allowed the construction of a concatenated phylogram of 45S fragments with most taxa. However, some specimens of the hexaploid *U. australis* Clemente showed two very divergent haplotypes in both fragments (in ITS and in ETS), and it was impossible to know which haplotypes of ITS and ETS belong to the same genome, making it impossible to concatenate. The same situation occurred with the 45S sequences of the hexaploid *U. europaeus* ssp. *europaeus,* obtained in the GenBank. Therefore, to analyse the phylogeny of these two taxa, we constructed two separated phylograms (non-concatenated) one for ETS and another for ITS.

To analyse the possibility of occurrence of Long Branch Attraction (LBA) in the 45S phylograms, we re-run the matrix of 45S sequences with the Bayesian and Maximum Parsimony approaches. We also calculated the mean for the uncorrected distances, between *U. micranthus,* suspected to be affected by LBA, and we graphed these distances using a multidimensional scaling, based on the uncorrected pairwise genetic distances.

## Results

Sequences from all taxa and studied populations, for 45S and 5S-IGS markers were obtained. The complete sequence data set consisted of 32 concatenated different sequences of ETS and ITS (i.e.: ITS2, 58S and ITS2), 32 different sequences of 5S-IGS, plus 45 and 31 sequences for the separated analysis of non-concatenated ETS and ITS fragments.

Overall, the Bayesian Inference approach resulted in better-solved and strongly supported phylogenetic trees. ETS and ITS resulted in a matrix with 459 and 574 characters, respectively. ETS showed higher variability and phylogenetic utility than ITS, with 44 parsimony informative characters, surpassing the 22 characters from ITS. The 5S-IGS spacer showed 38 parsimony informative characters, although it was the smallest fragment, with only 248 base pairs. Many plants shared the same haplotypes, or very similar ones, even belonging to different taxa. Phylograms are depicted in figures 2 to 5.

**Fig 1.**
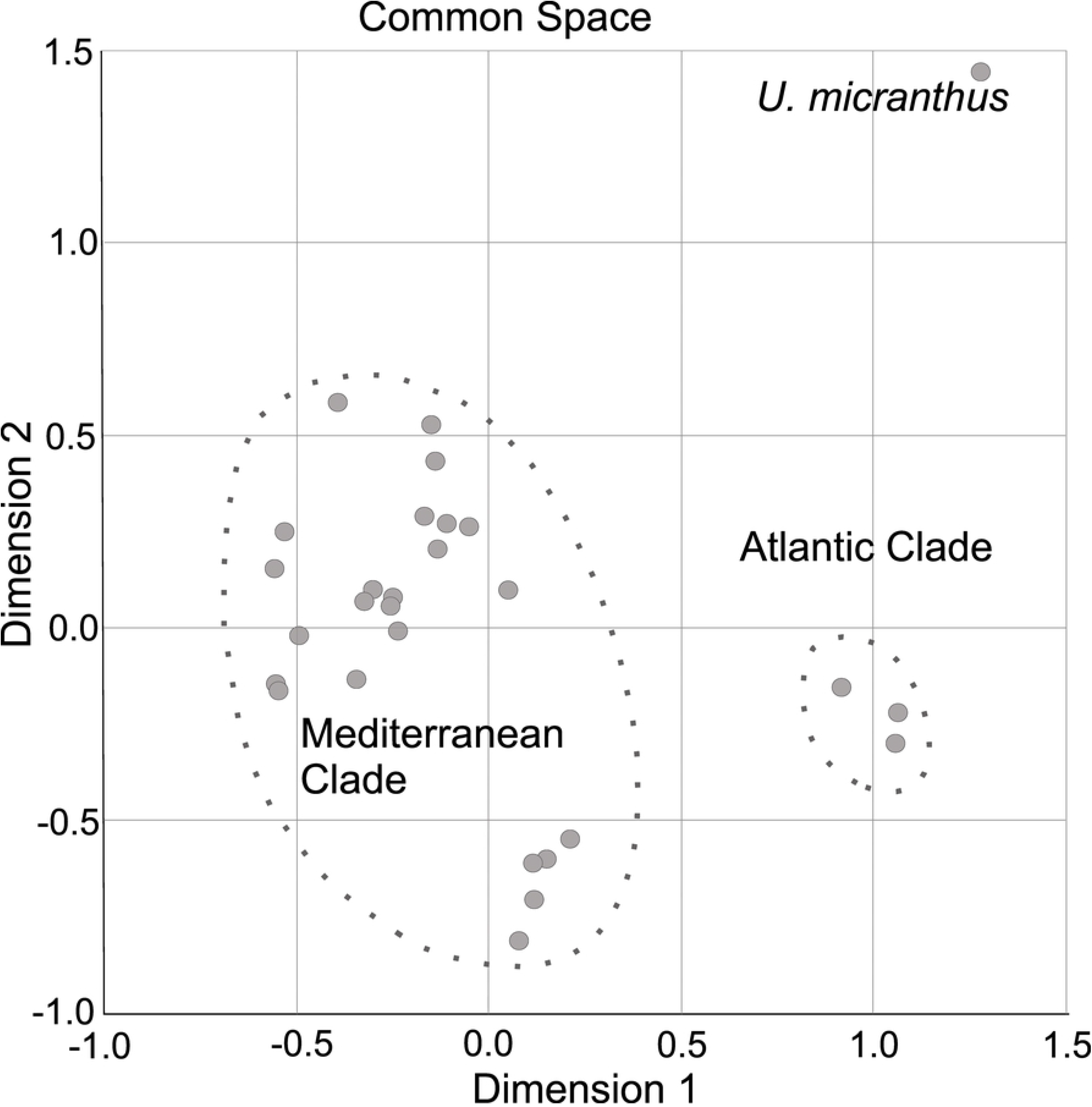
Multidimensional scaling, based on the uncorrected pairwise genetic distances (stress=0.148) of 45S haplotypes.

**Fig 2.**
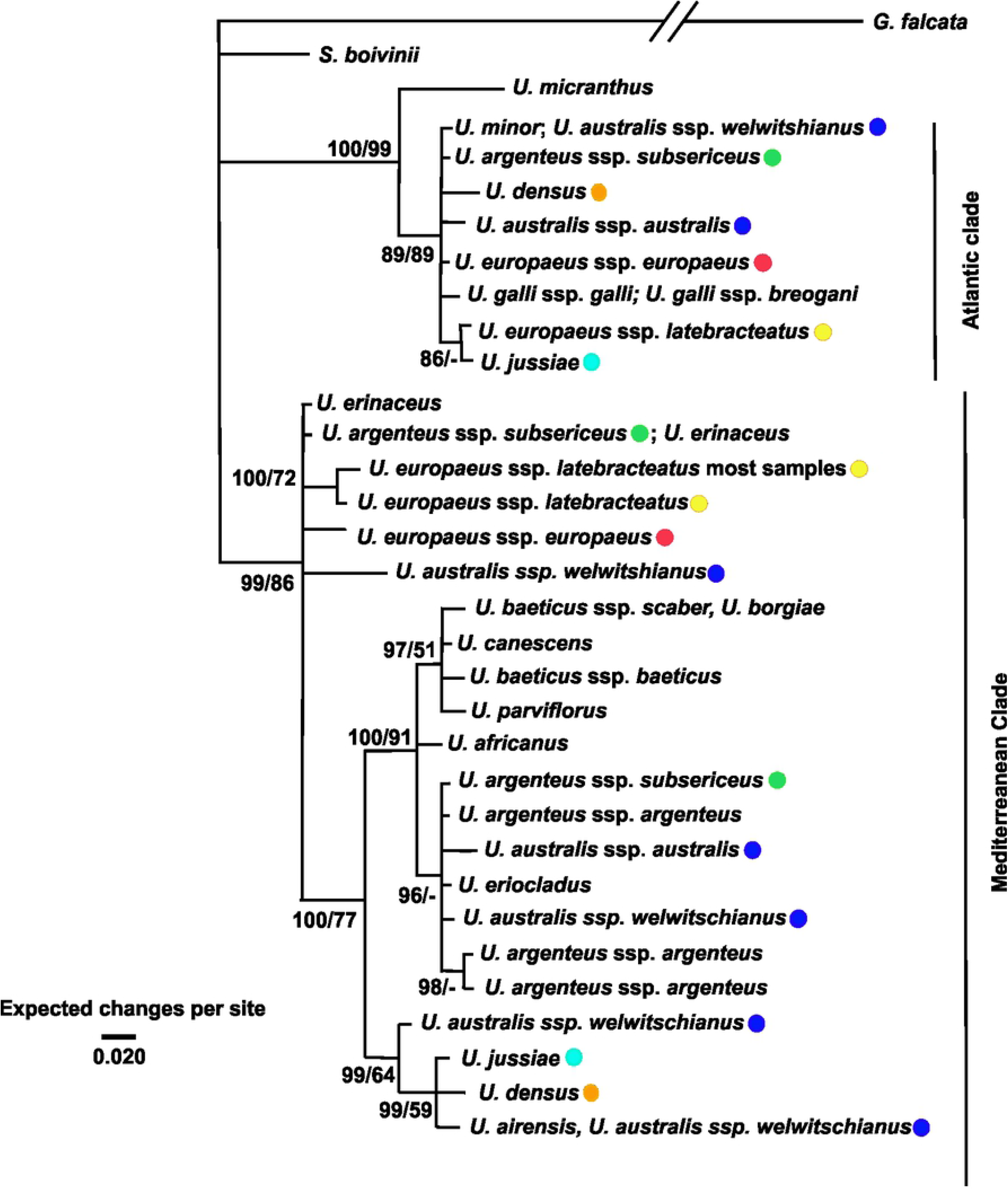
Fifty percent Bayesian majority-rule consensus tree from Bayesian inference of 5S-Intergenic Spacer. Posterior probabilities and bootstrap support values for Maximum Parsimony for each node were separated by a bar, in the left and in the right, respectively.

**Fig 3.**
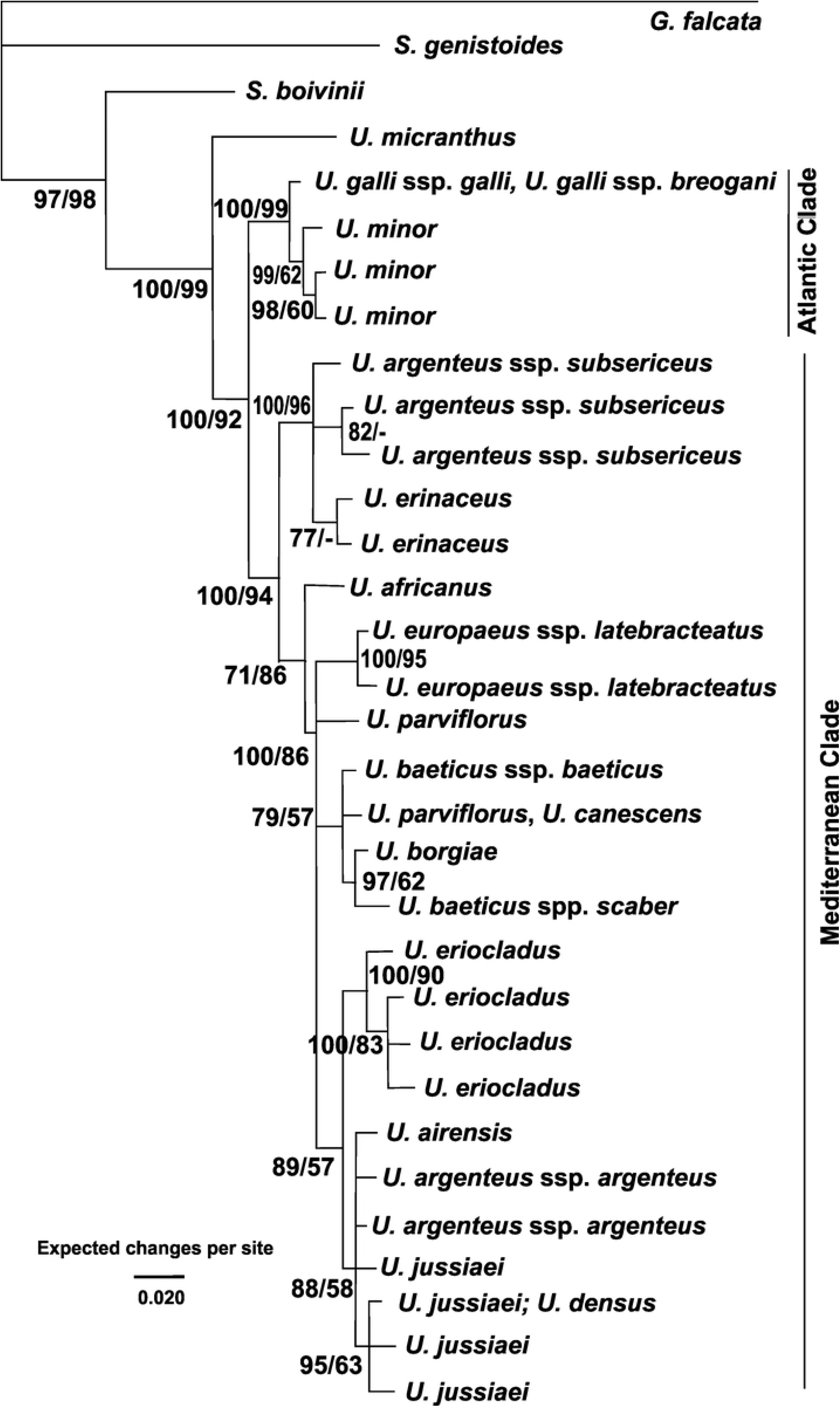
Fifty percent Bayesian majority-rule consensus tree from Bayesian inference of concatenated 45 fragments (ETS, ITS1, 5.8S and ITS2). Nodes supported by less than 0.60 of posterior probabilities were collapsed. Posterior probabilities and bootstrap support values higher than 50 for Maximum Parsimony, for each node were separated by a bar, in the left and in the right, respectively.

**Fig 4.**
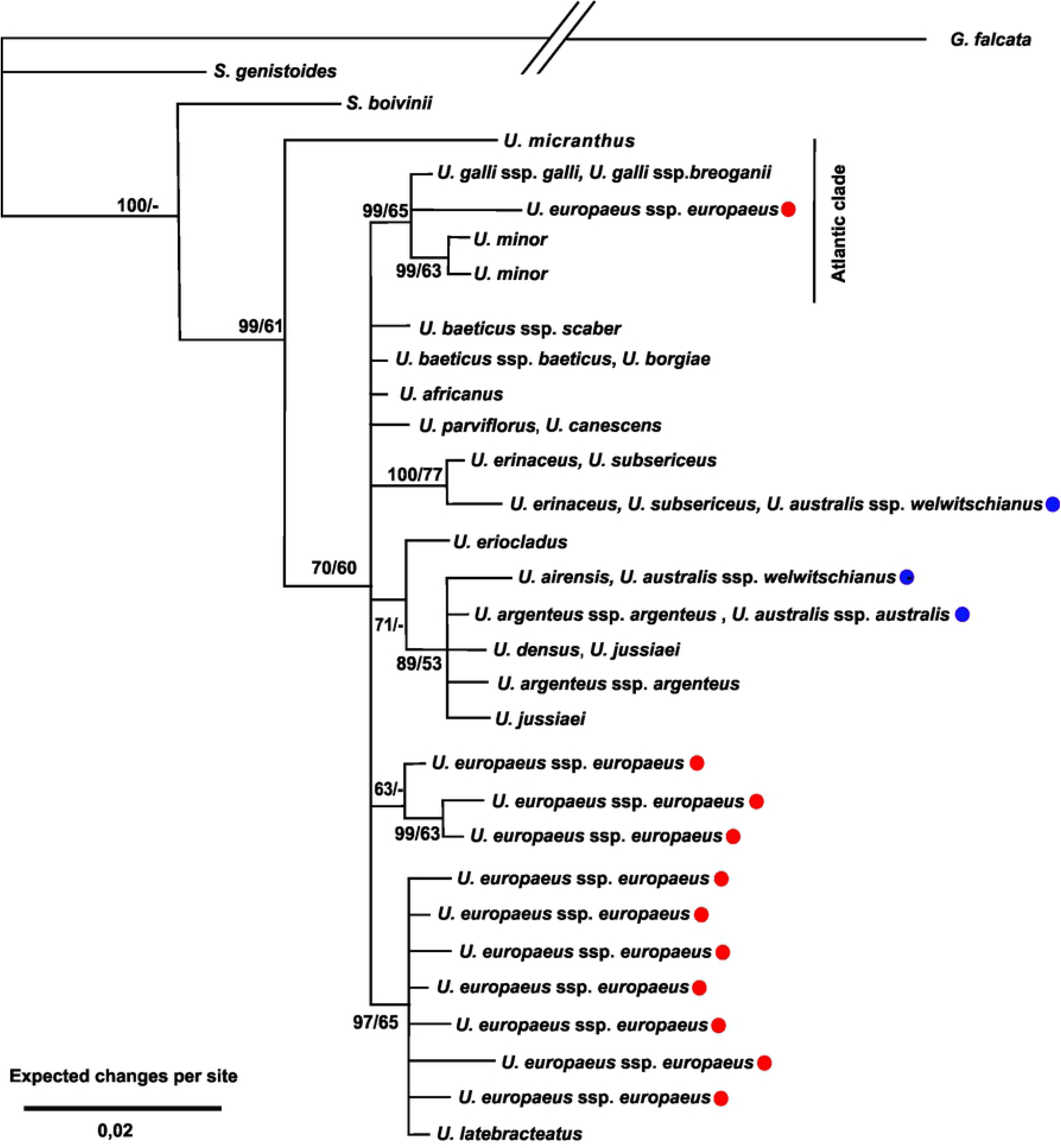
Fifty percent Bayesian majority-rule consensus tree from Bayesian inference of the ITS fragment (ITS1, 5.8S and ITS2), with the sequences of the polyploids *U. europaeus* ssp. *europaeus* and *U. australis*. Nodes supported by less than 0.60 of posterior probabilities were collapsed. Posterior probabilities and bootstrap support values higher than 50 for Maximum Parsimony for each node were separated by a bar, in the left and in the right, respectively.

**Fig 5.**
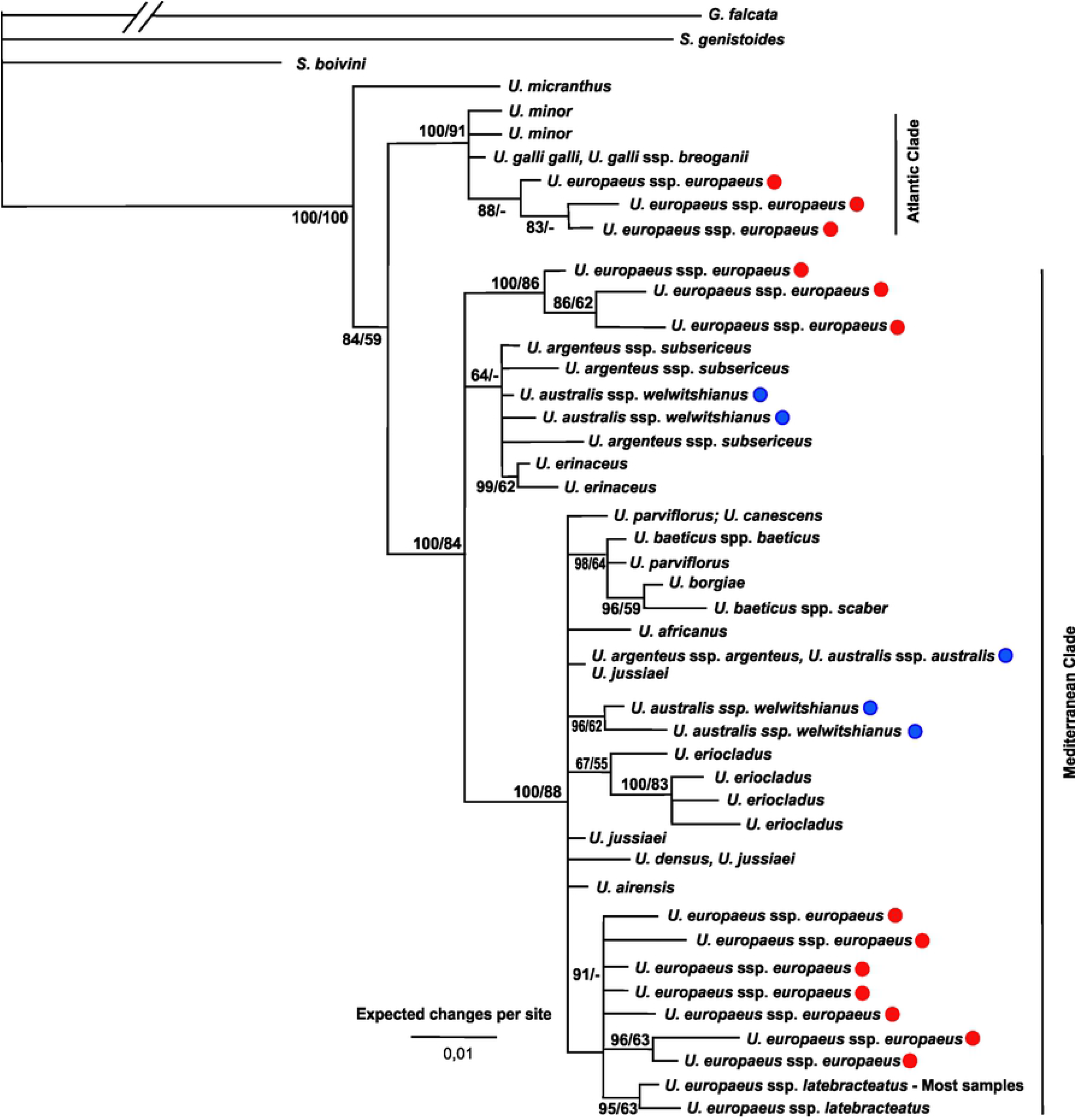
Fifty percent Bayesian majority-rule consensus tree from Bayesian inference of the external transcribed spacer, with the sequences of the polyploids *U. europaeus* ssp. *europaeus* and *U. australis*. Nodes supported by less than 0.60 of posterior probabilities were collapsed. Posterior probabilities and bootstrap support values higher than 50 for Maximum Parsimony for each node were separated by a bar, in the left and in the right, respectively.

All diploid taxa showed only one sequence of 5S-IGS and 45S, or two very similar sequences, with a maximum of two double peaks, even when the fragments were amplified with non-allelespecific sequencing primers. When tested with different allele-specific sequencing primers for 5S-IGS haplotypes, diploids did not reveal different haplotypes, independently of the primer used.

The polyploids *U. gallii* Planch. ssp. *gallii, U. gallii* Planch ssp. *breoganii* (Castroviejo & Valdes-Bermejo) Rivas Mart. et al., *U. eriocladus* Vicioso, *U. erinaceus* Welw. Ex Webb. and *U. borgiae* showed only one, or few very similar haplotypes, distinguished by few mutations, which grouped in the same clades, in both markers (45S and 5S-IGS).

Contrariwise, the haplotypes of the polyploids *U. australis*, *U. densus* Welw. ex Webb., *U. jussiaei* Webb., *U. argenteus* ssp. *subsericeus, U. europaeus* L. ssp. *latebracteatus* (Mariz) Rothm. and *U. europaeus* ssp. *europaeus* showed strong polymorphism in 5S-IGS and, at least, two different haplotypes grouped in very distant positions in the phylogram. Contrasting with their diversity in 5S-IGS, these polyploids showed low diversity in the 45S, with two exceptions, and all haplotypes from the same taxa grouped together in the same clades. Those exceptions were *Ulex europaeus* subsp. *europaeus* and *U. australis,* which also showed a strong polymorphism in both 45S and 5S-IGS haplotypes.

Regarding only the diploid *taxa* and the polyploids with low haplotype diversity, we can identify two main clades, in both 5S-IGS and 45S markers. One grouped the species which occur mainly in Atlantic bioclimate or in wet soils (henceforth called Atlantic Clade), namely *U. minor* Roth., *U. gallii* ssp. *breoganii, U. gallii* ssp. *gallii* and, in the 5S-IGS phylogram, *U*. *micranthus*. The other clade grouped all the remaining diploids and polyploids with low haplotype diversity that occur mainly in Mediterranean bioclimate (henceforth called Mediterranean Clade). The polyploid *taxa* with high diversity of haplotypes showed haplotypes clustered in both clades.

*Ulex australis, U. densus* Welw. ex Webb., *U. jussiaei* Webb., *U. argenteus* ssp. *subsericeus, U. europaeus* L. ssp. *latebracteatus* (Mariz) Rothm. and *U. europaeus* ssp. *europaeus* showed haplotypes (in 5S-IGS and/or 45S) clustered in the Atlantic Clade and in the Mediterranean Clade.

*Ulex europaeus* ssp. *europaeus* showed haplotypes of 45S grouped in three different positions (Figures 4 and 5), although presenting only two different haplotypes in 5S-IGS. *Ulex argenteus* ssp. *subsericeus* showed three haplotypes in 5S-IGS phylogram (Figure 2): one shared with *U. erinaceus,* grouping as an early branching of Mediterranean Clade, a second clustered with *U. argenteus* Welw. Ex Webb ssp*. argenteus,* and a third clustered in the Atlantic Clade (Figure 2). All 45S haplotypes of this taxon grouped together with *U. erinaceus,* in a subclade, which is early branching the Mediterranean Clade (Figure 3).

Results from the hexaploid *U. australis* are complex and showed a high diversity of haplotypes. We emphasize five points: 1) all plants from the same populations showed the same or very similar haplotypes; 2) in both markers (45S and 5S-IGS), *U. australis* showed haplotypes which grouped in different clades; 3) the haplotypes of different populations grouped in different positions in the phylograms of 5S-IGS, ETS and ITS (Figures 2, 4 and 5); 4) two from the five studied populations showed two different haplotypes of 45S, while the other three populations showed just one 45S haplotype; 5) all populations showed different combinations of haplotypes, grouping in different positions.

The 45S haplotypes of the hexaploid *U. jussiaei* and the tetraploid *U. densus* grouped together in the same clade with *U. airensis* Espírito-Santo, Cubas, Lousã, Pardo, & Costa. In 5S-IGS, both taxa showed two haplotypes, one of which also grouped together with *U. airensis*, within the Mediterranean Clade, and the other in the Atlantic Clade (Figures 2 and 3).

In the 5S-IGS phylogram (Figure 2), *U. europaeus* ssp. *latebracteatus,* showed two haplotypes, one nested in the Atlantic Clade and the other grouped as an early branching of the Mediterranean Clade. In the 45S, it grouped also within the Mediterranean Clade, not as an early branching, but in a subclade with most diploids.

Within the Mediterranean Clade, there is one subclade which group the diploids *U. airensis*, *U. parviflorus* Pourr.*, U. africanus* Webb, *U. argenteus* ssp. *argenteus, U. baeticus* Boiss., *U. canescens* Lange, and some polyploids. This subclade is well supported in both 45S and 5S-IGS phylograms. *Ulex airensis* is a species separated from *U. parviflorus* in 1997 (33) and groups in different positions from *U. parviflorus,* in both the 45S and 5S-IGS phylograms.

The position of the diploid *U. argenteus* is incongruent, as it grouped with *U. airensis* in the 45S phylogram (Figure 3, although this subclade is not well supported: Bayesian posterior probabilities and bootstrap are 89 and 57, respectively), but grouped with *U. eriocladus* in 5S-IGS phylogram (Figure 2).

The phylograms of 5S-IGS and 45S showed an incongruence in the position of *U. micranthus*. This species grouped as the first divergence of the genus in the 45S phylograms, but in 5S, grouped as a sister species of the Atlantic Clade. As stated before, we rerun the Bayesian and the Parsimony analyses without outgroups of the 45S matrix and found that *U. micranthus* grouped as an early branching of the Atlantic Clade, as happened in 5S-IGS phylogram (trees not shown). Furthermore, the mean of the uncorrected distance between *U. micranthus* and the Atlantic Clade (mean distance 2,486 + 0,049 %) is lower than the distance to the Mediterranean Clade (mean distance 2,699+0,001), and also lower than the mean between *U. micranthus* and all other taxa (mean distance 2,669+0,001), reinforcing the hypothesis of a closer phylogenetic relationship between *U. micranthus* and the Atlantic Clade. The figure 1 summarizes the relationships among species, using a multidimensional scaling, based on the uncorrected pairwise genetic distances (stress=0.148). The inspection of this figure corroborates the previous analysis and reveals that the 45S haplotypes of *U. micranthus* are closer to the Atlantic Clade, as expected.

## Discussion

### Evidence of reticulated evolution

As previously mentioned, the data from nrDNA may prove informative to documenting the historical hybridization events, providing information on paternal lineages (10). Thus, since the early 90s, the presence of very divergent nrDNA haplotypes in polyploids is interpreted as an evidence of a hybrid origin, indicating that those taxa are allopolyploids (34,35). According to this, in what concerns our data, the genus *Ulex* includes, at least, seven allopolyploids: *U. australis* ssp. *australis* Clemente, *U. australis* Clemente ssp. *welwitschianus* (Planch.) Esp. Santo et al., *U. densus, U. jussiaei, U. argenteus* ssp. *subsericeus, U. europaeus* ssp. *europaeus* and *U. europaeus* ssp. *latebracteatus,* because all of them showed two or more very different haplotypes in 5S-IGS. *U. australis* ssp. *welwitschianus* and *U. europaeus* ssp. *europaeus* also showed divergent haplotypes in 45S.

It is interesting to note that most of these polyploids register a higher haplotype diversity in 5S-IGS than in 45S. Some previous studies reported that, in 5S, the concerted evolution leads to the homogenization intra-loci, but does not act, or does it very weakly, inter-loci (3,8,9). Based on similar results, Mahelka et al. (2013) advocate that 5S nrDNA may provide a more suitable marker for reconstructing the histories of allopolyploid species than 45S. Therefore, the higher intraspecific diversity of 5S-IGS in *Ulex* polyploids could be interpreted as a consequence of the higher effectiveness of inter-loci concerted evolution in 45S and the slowness, or absence, of inter-loci concerted evolution in 5S.

*Ulex europaeus* ssp. *europaeus* showed also very divergent haplotypes in 45S and 5S-IGS (Figures 2, 4 and 5). We interpret this result as a confirmation of allohexaploidy, as stated previously by Dias et al.(20), based only in the 45S markers. However, differently from the other polyploids, the 45S haplotypes were more diverse (ITS and ETS phylograms, figures 4 and 5), grouping in three main locations, while 5S-IGS haplotypes just groups twice (Figure 2).

Beside concerted evolution, loss of loci may also have contributed to the absence of one parental 5S-IGS haplotypes, as well as 45S haplotypes, in allopolyploids. Furthermore, some authors suggested that this phenomenon is more likely to occur in higher ploidy levels (36,37). This could explain the absence of a third haplotype of 5S-IGS in hexaploid *U. europaeus* ssp. *europaeus.* Nevertheless, other hypotheses could be considered, including the mismatch of the allele-specific sequencing primers used to amplify 5S-IGS haplotypes, which may have prevented the amplification of a third 5S-IGS haplotype in *U. europaeus* ssp. *europaeus*.

Regarding the identification of parental lineages, we emphasize that *U. europaeus* ssp. *europaeus* presented phylogenetic connections with three different lineages (Figures 2, 4 and 5): 1) the lineage of *U. minor* (Atlantic Clade), connection supported by the data from 5S-IGS, ITS and ETS; 2) an early branching within the Mediterranean Clade, also supported by the data from 5S-IGS and ETS; 3) and a third lineage belonging to a subclade of Mediterranean Clade, supported only by ETS phylogram.

The hexaploid *U. argenteus* ssp. *subsericeus* showed three very similar 45S haplotypes, but also three very different in 5S-IGS. Both markers indicate genetic connection with *U. erinaceus* because these taxa grouped together in the 45S phylogram in a well-supported clade and shared one 5S-IGS haplotype. The other two 5S-IGS haplotypes from *U. argenteus* ssp. *subsericeus* grouped in two different clades: together with the diploids *U. argenteus* ssp. *argenteus;* and together with *U. minor.* These data suggest that it is an allohexaploid originated from these three lineages. Nowadays, *U. argenteus* ssp. *argenteus, U. minor* and *U. erinaceus* are not sympatric, but they occur in the extreme south of Iberian Peninsula, which suggests that they may have been in contact in a not-so-distant past.

Regarding *U. australis,* the presence of haplotypes of 5S-IGS, ITS and ETS, which grouped in three different positions in the phylograms, suggests that this species is an allohexaploid. However, each population was represented by a different combination of haplotypes, which can be explained, at least, in two ways: 1) the concerted evolution, or the loss of loci, fixed different haplotypes in different populations, or 2) under the designation of *U. australis* there are different unknown taxa with different phylogenetic origins. The former hypothesis is noteworthy because there are few reports on the literature of bidirectional concerted evolution of rnDNA in different populations of the same species (38,39). The second hypothesis seems unlikely because if each the combination of haplotypes represents a different taxon, there would be, at least, five different taxa, with no obvious morphological or ecological differences. However, an extensive sample and a detailed analysis of genetic markers and morphological characters through the geographical range of *U. australis* is required to fully clarify this question.

The tetraploid *U. densus* and the hexaploid *U. jussiaei* showed similar phylogenetic connections with two lineages: the lineage of the diploid *U. minor* (the Atlantic Clade), supported by one haplotype of 5S-IGS, and with a subclade of the Mediterranean Clade, together with the diploid *U. airensis,* supported by a second haplotype of 5S-IGS and by 45S. These results suggest that they are allopolyploids, sharing these two parental diploid lineages. Despite the similarity of haplotypes, these taxa have deep morphological and ecological differences (18). These differences could arise from a hypothetical third genome involved in the origin of the hexaploid *U. jussiaei,* although not detected in our analysis, from different genetic contributions from these two putative parental lineages, or from evolution subsequent to the hybridization events.

The phylogeny of the tetraploid *U. europaeus* ssp. *latebracteatus* is harder to clarify with our data. In the 5S-IGS phylogram, three haplotypes grouped in two distant positions: one in the Atlantic Clade and two as an early branching of the Mediterranean Clade. These results suggest that *U. europaeus* ssp*. latebracteatus* is an allotetraploid. However, there is an incongruence: in the 45S phylogram, it grouped within the Mediterranean Clade, although not as an early branching (Figure 3). Despite this, *U. europaeus* ssp. *latebracteatus* grouped together with *U. europaeus* ssp. *europaeus* in the phylograms of 5S-IGS, ITS and ETS (Figures 2, 4 and 5), which suggests a close phylogenetic connection.

If our hypothesis is correct, the tetraploid *U. europaeus* ssp. *latebracteatus* arose from only two different genomes, while the hexaploidy of *U. europaeus* ssp. *europaeus* resulted from three different genomes, which means that these two taxa have different phylogenetic origins. Furthermore, Cubas & Pardo (40) showed that *U. europaeus* ssp. *europaeus* and *U. europaeus* ssp. *latebracteatus* have distinct morphological characters, which are maintained in sympatric populations, and argued that ploidy acts as an effective reproductive barrier (40). Therefore, they do not fit the biological nor the phylogenetic species concepts and should be assign as different species.

### Other evolutionary insights

Five polyploids, namely: *U. gallii* ssp. *gallii, U. gallii* ssp. *breoganii, U. eriocladus, U. erinaceus* and *U. borgiae,* display low intraspecific diversity both in 45S and 5S-IGS haplotypes, and they do not show topological incongruences between the 45S and 5S-IGS phylograms. They can be autopolyploids, or they can be allopolyploids in which the homogenization process of nrDNA is completed, and the alleles of 45S and 5S-IGS from same parental species were lost. Therefore, the autopolyploid origin of these taxa needs confirmation by additional genetic markers. *Ulex borgiae* is probably an example of this, as Cubas et al. (18), using microsatellite (CpSSR) markers, suggested that it arose through hybridization of *U. baeticus* ssp. *baeticus* and *U. baeticus* ssp. *scaber*, two very close taxa.

If these species are autopolyploids, the hexaploid *U. gallii* ssp. *gallii* and the tetraploid *U. gallii* ssp. *breoganii* could compose a polyploid series with the diploid *U. minor.* We stress that the distinctive morphological characters that separate these three species are biometric (18), as usual in taxa that diverge by ploidy. Lastly, if the tetraploids *U. erinaceus* and *U. eriocladus* are autopolyploids we did not find the correspondent diploids.

Concerning diploids, we emphasize that the Mediterranean Clade grouped seven diploid taxa, namely *U. argenteus, U. africanus, U. airensis, U. baeticus* ssp. *baeticus, U. baeticus* ssp. *scaber, U. canescens* and *U. parviflorus,* all occurring in allopatry (Figure 6), suggesting that the speciation of diploid lineages, in *Ulex,* occurred mainly by allopatric or parapatric speciation.

**Fig 6.**
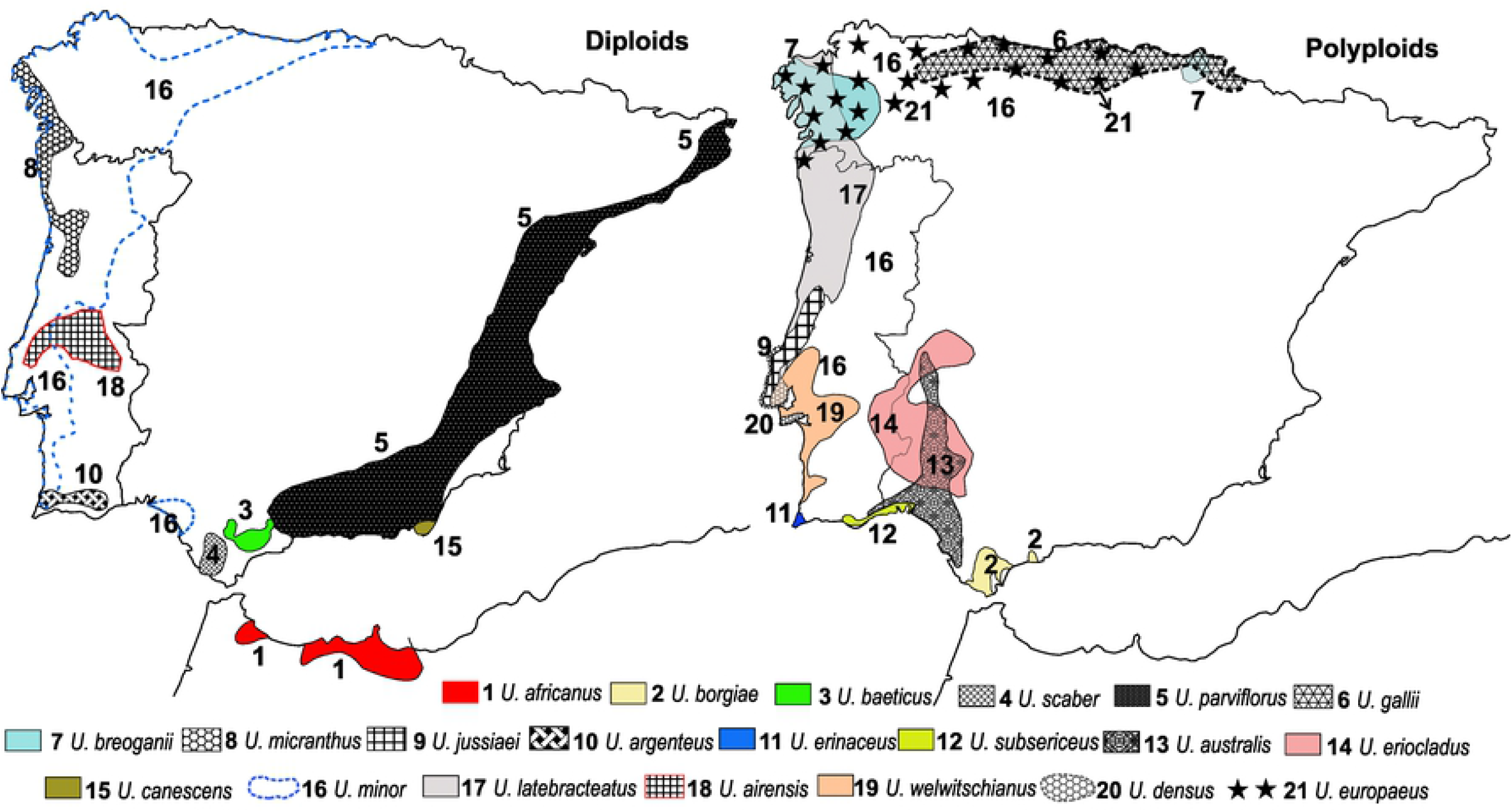
Geographical distribution of studied *taxa*, based in Anthos (n.d.) Cubas et al., (2005) and; Flora-On (2014)

*Ulex parviflorus,* both subspecies of *U. baeticus,* and *U. canescens* grouped in the same subclade, within the Mediterranean Clade, in both 5S-IGS and 45S phylograms. These results support a close phylogenetic relationship between these three taxa.

As stated before, in the phylograms of both markers, *U. erinaceus* groups as an early branching of Mediterranean Clade, not in this subclade of *U. canescens*. Accordingly, the morphological similarity between both stenoendemisms: *U. canescens* from Cape of Gata (Spain) and *U. erinaceus,* endemic from Cape of Sagres (Portugal), namely the presence of a dense branching and a cushion-like form (18), is probably a homoplasy, resulting from exposure to strong winds and dry habitats.

The incongruence between the positions of *U. micranthus* in the trees of 5S and 45S could result from Long Branch Attraction. This is a known methodological bias which leads to the erroneous grouping of two or more long branches as sister groups, in which a ingroup taxa can be ‘pulled’ to the base of the tree by long-branched outgroups (41,42). As far as we know, there is no method to prove that a particular topology results from Long Branch Attraction. Bergsten (2005) argued that the reconstruction of the trees without outgroups (unrooted trees) can indicate the corrected position of the taxon suspected of erroneous grouping. As pointed by Borowiec et al. (42), if the outgroups were to influence the branching order, we would expect the internal topology to be changed, if they were excluded. Indeed, the construction of unrooted trees of Parsimony and Bayesian approaches confirmed that *U. micranthus* grouped within the Atlantic Clade as an early branching, as it happened in the phylogram of 5S-IGS. Furthermore, the uncorrected distance between the *U. micranthus* and the Atlantic Clade is lower than the distance to the Mediterranean Clade, as well between *U. micranthus* and all other taxa. Future studies using data from other markers are needed to corroborate this hypothesis.

This is an interesting hypothesis, because, if confirmed, the first split within the genus *Ulex* would divide hydrophytes and xerophytes diploids. Allopolyploid *taxa* seem to arise from hybridation of these hydrophytes and xerophytes diploid lineages. We emphasize that *U. micranthus, U. gallii* ssp. *gallii* and *U. gallii* ssp. *breoganii* occur almost exclusively in the Atlantic Bioregion, although *U. minor* exists in both Atlantic and Mediterranean Bioregions. Nevertheless, under Mediterranean climate, *U. minor* always occurs in marshes or moist soils, usually with edaphic water compensation.

### Taxonomic consequences

Our data can help to clarify some issues in *Ulex* taxonomy. Allopolyploidy provides a high degree of reproductive isolation from other ploidies, even with the progenitor species (43). For this reason, allopolyploids should not be assigned as subspecies, once there are not conspecific with lower ploidies, and should be assigned as species. This is the case of *U. argenteus* ssp. *subsericeus*, *U. europaeus* ssp. *europaeus* and *U. europaeus* ssp. *latebracteatus*.

Both markers (5S-IGS and 45S) confirm the splitting between *Ulex airensis* and *U. parviflorus,* as proposed by Espírito-Santo et al (33), and the validity of *Ulex airensis* as a different species. Lastly, Rothmaler (44) synonymized the stenoendemisms *U. canescens* and *U. erinaceus*, being followed by Guinea & Webb (16). Our results showed *U. canescens* is phylogenetically closer to *U. parviflorus* and *U. baeticus,* supporting the splitting between both stenoendemisms.

## Conclusions

The main conclusion of this study is the confirmation of the hypothesis brought by Ainouche et al. (2003), that *Ulex* radiation occurred mainly by polyploidization, especially allopolyploidy. Indeed, allopolyploidy, as a speciation mechanism, is supported by ribosomal DNA markers, explaining the origin of, at least, seven taxa: the hexaploids *U. australis* ssp. *australis, U. australis* ssp. *welwitschianus, U. argenteus* ssp. *subsericeus, U. jussiaei, U. europaeus* ssp. *europaeus,* and the tetraploids *U. densus* and *U. europaeus* ssp. *latebracteatus.* This number is a third of the 21 studied taxa. However, our data are less informative on the origin of the polyploids *U. gallii* ssp. *gallii, U. gallii* ssp. *breoganii, U. eriocladus, U. erinaceus* and *U. borgiae,* which can be allopolyploids or autopolyploids. This approach should be complemented with additional genetic markers. This set of new data have important consequences in the taxonomy of *Ulex,* namely the assignment of specific rank to the allopolyploids, the confirmation of the validity of *U. airensis* as a species, and the splitting between *U. canescens* and *U. erinaceus*.

Our data also suggests, although do not prove, that the first splitting in this genus separate xerophytic and hydrophilic diploid. All the putative allopolyploids seem to be originated from the hybridization of these two early diverging lineages.

Other important result is the confirmation, for the genus *Ulex,* that 5S-IGS provided a more suitable marker for reconstructing histories of allopolyploid species than 45S, as stated by Mahelka et al. (3), probably because the inter-loci homogenization is less effective in 5S than in 45S repeats.

## Acknowledgments

We thank to Francisco Javier Amigo Vazquez for the supply of samples of *Ulex gallii* ssp. *breoganii* and to Carlos Vila-Viçosa for the suggestions and very careful review of our paper.

## References

1. Wendel JF, Jackson SA, Meyers BC, Wing RA. Evolution of plant genome architecture. Genome Biol [Internet]. 2016 Dec 1;17(1):37. Available from: https://genomebiology.biomedcentral.com/articles/10.1186/s13059-016-0908-1

2. Wicke S, Costa A, Muñoz J, Quandt D. Restless 5S: The re-arrangement(s) and evolution of the nuclear ribosomal DNA in land plants. Mol Phylogenet Evol [Internet]. 2011 Nov;61(2):321–32. Available from: https://linkinghub.elsevier.com/retrieve/pii/S1055790311003125

3. Mahelka V, Kopecký D, Baum BR. Contrasting Patterns of Evolution of 45S and 5S rDNA Families Uncover New Aspects in the Genome Constitution of the Agronomically Important Grass Thinopyrum intermedium (Triticeae). Mol Biol Evol [Internet]. 2013 Sep;30(9):2065–86. Available from: https://academic.oup.com/mbe/article-lookup/doi/10.1093/molbev/mst106

4. Poczai P, Hyvönen J. Nuclear ribosomal spacer regions in plant phylogenetics: problems and prospects. Mol Biol Rep [Internet]. 2010 Apr 21;37(4):1897–912. Available from: http://link.springer.com/10.1007/s11033-009-9630-3

5. Sastri DC, Hilu K, Appels R, Lagudah ES, Playford J, Baum BR. An overview of evolution in plant 5S DNA. Plant Syst Evol [Internet]. 1992;183(3-4):169–81. Available from: http://link.springer.com/10.1007/BF00940801

6. Baum BR, Feldman M. Elimination of 5S DNA unit classes in newly formed allopolyploids of the genera Aegilops and Triticum. Genome [Internet]. 2010 Jun;53(6):430–8. Available from: http://www.nrcresearchpress.com/doi/10.1139/G10-017

7. Kotseruba V, Pistrick K, Blattner FR, Kumke K, Weiss O, Rutten T, et al. The evolution of the hexaploid grass Zingeria kochii (Mez) Tzvel. (2n=12) was accompanied by complex hybridization and uniparental loss of ribosomal DNA. Mol Phylogenet Evol [Internet]. 2010 Jul;56(1):146–55. Available from: https://linkinghub.elsevier.com/retrieve/pii/S1055790310000059

8. Cronn RC, Zhao X, Paterson AH, Wendell JF. Polymorphism and concerted evolution in a tandemly repeated gene family: 5S ribosomal DNA in diploid and allopolyploid cottons. J Mol Evol [Internet]. 1996 Jun;42(6):685–705. Available from: http://link.springer.com/10.1007/BF02338802

9. Fulnecěk J, Lim KY, Leitch AR, Kovarĭk A, Matyásěk R. Evolution and structure of 5S rDNA loci in allotetraploid Nicotiana tabacum and its putative parental species. Heredity (Edinb) [Internet]. 2002 Jan 24;88(1):19–25. Available from: http://www.nature.com/articles/6800001

10. Álvarez I, Wendel JF. Ribosomal ITS sequences and plant phylogenetic inference. Mol Phylogenet Evol [Internet]. 2003 Dec;29(3):417–34. Available from: https://linkinghub.elsevier.com/retrieve/pii/S1055790303002082

11. Sang T, Crawford DJ, Stuessy TF. Documentation of reticulate evolution in peonies (Paeonia) using internal transcribed spacer sequences of nuclear ribosomal DNA: implications for biogeography and concerted evolution. Proc Natl Acad Sci [Internet]. 1995 Jul 18;92(15):6813–7. Available from: http://www.pnas.org/cgi/doi/10.1073/pnas.92.15.6813

12. Liu Z-L, Zhang D, Wang X-Q, Ma X-F, Wang X-R. Intragenomic and interspecific 5S rDNA sequence variation in five Asian pines. Am J Bot [Internet]. 2003 Jan 1;90(1):17–24. Available from: http://doi.wiley.com/10.3732/ajb.90.1.17

13. Bao Y, Wendel JF, Ge S. Multiple patterns of rDNA evolution following polyploidy in Oryza. Mol Phylogenet Evol [Internet]. 2010 Apr;55(1):136–42. Available from: https://linkinghub.elsevier.com/retrieve/pii/S1055790309004175

14. Ainouche A, Bayer RJ, Cubas P, Misset MT. Phylogenetic relationships within tribe Genisteae (Papilionoideae) with special reference to genus Ulex. In: Klitgaard B., Bruneau A, editors. Advances in Legume Systematics part 10, Higher Level Systematics. Kew: Royal Botanic Gardens; 2003. p. 239–52.

15. Cubas P, Pardo C, Tahiri H. Genetic variation and relationships among Ulex (Fabaceae) species in southern Spain and northern Morocco assessed by chloroplast microsatellite (cpSSR) markers. Am J Bot [Internet]. 2005 Dec 1;92(12):2031–43. Available from: http://doi.wiley.com/10.3732/ajb.92.12.2031

16. Guinea E, Webb DA. Ulex. In:Tutin TG, editor. Flora Europaea. Cambridge: Cambridge University Press.; 1968. p. 102–3.

17. Maire R. Dicots. Leguminosae, part. In:Quezel P, editor. Flore de l’Afrique du Nord. 1987.

18. Cubas P. Ulex L. In:Talavera S, Aedo C, Castroviejo S, Romero Zarco C, Saez L, Salgueiro FJ, et al., editors. Flora iberica. Madrid: CSIC; 2000. p. 212–239.

19. Pardo C, Cubas P, Tahiri H. Molecular phylogeny and systematics of Genista (Leguminosae) and related genera based on nucleotide sequences of nrDNA (ITS region) and cpDNA (trn L-trn F intergenic spacer). Plant Syst Evol [Internet]. 2004 Feb 1;244(1-2):93–119. Available from: http://link.springer.com/10.1007/s00606-003-0091-1

20. Dias PB, Bellot S, Affagard M, Aïnouche ML, Misset MT, Aïnouche A. On the allopolyploid origin of the invasive European gorses, Ulex europaeus subsp. Europaeus (Fabaceae; Genisteae). In: French Annual Polyploidy and Cytogenetics Meeting. Rennes; 2013.

21. Sun Y, Fung K-P, Leung P-C, Shaw P-C. A phylogenetic analysis of Epimedium (Berberidaceae) based on nuclear ribosomal DNA sequences. Mol Phylogenet Evol [Internet]. 2005 Apr;35(1):287–91. Available from: https://linkinghub.elsevier.com/retrieve/pii/S1055790304004038

22. Rauscher JT, Doyle JJ, Brown AHD. Internal transcribed spacer repeat-specific primers and the analysis of hybridization in the Glycine tomentella (Leguminosae) polyploid complex. Mol Ecol [Internet]. 2002 Dec;11(12):2691–702. Available from: http://doi.wiley.com/10.1046/j.1365-294X.2002.01640.x

23. Scheen A-C, Pfeil BE, Petri A, Heidari N, Nylinder S, Oxelman B. Use of allele-specific sequencing primers is an efficient alternative to PCR subcloning of low-copy nuclear genes. Mol Ecol Resour [Internet]. 2012 Jan;12(1):128–35. Available from: http://doi.wiley.com/10.1111/j.1755-0998.2011.03070.x

24. Sousa-Santos C, Robalo JI, Collares-Pereira M-J, Almada VC. Heterozygous indels as useful tools in the reconstruction of DNA sequences and in the assessment of ploidy level and genomic constitution of hybrid organisms. DNA Seq [Internet]. 2005 Jan 11;16(6):462–7. Available from: http://www.tandfonline.com/doi/full/10.1080/10425170500356065

25. Hall TA. BioEdit (v. 7.0.1): a user-friendly biological sequence alignment editor and analysis program for Windows 95/98/NT. Nucleic Acids Symp Ser. 1999;41:95–98.

26. Thompson J, Gibson TJ, Plewniak F, Jeanmougin F, Higgins DG. The CLUSTAL_X windows interface: flexible strategies for multiple sequence alignment aided by quality analysis tools. Nucleic Acids Res [Internet]. 1997 Dec 15;25(24):4876–82. Available from: https://academic.oup.com/nar/article-lookup/doi/10.1093/nar/25.24.4876

27. Ronquist F, Huelsenbeck JP. MrBayes 3: Bayesian phylogenetic inference under mixed models. Bioinformatics [Internet]. 2003 Aug 12;19(12):1572–4. Available from: https://academic.oup.com/bioinformatics/article-lookup/doi/10.1093/bioinformatics/btg180

28. Swofford DL. PAUP*. Phylogenetic Analysis Using Parsimony (*and Other Methods). Version 4 beta 10. Sunderland: Sinauer Associates; 2003.

29. Felsenstein J. CONFIDENCE LIMITS ON PHYLOGENIES: AN APPROACH USING THE BOOTSTRAP. Evolution (N Y) [Internet]. 1985 Jul;39(4):783–91. Available from: http://doi.wiley.com/10.1111/j.1558-5646.1985.tb00420.x

30. Posada D. jModelTest: Phylogenetic Model Averaging. Mol Biol Evol [Internet]. 2008 Apr 3;25(7):1253–6. Available from: https://academic.oup.com/mbe/article-lookup/doi/10.1093/molbev/msn083

31. Guindon S, Gascuel O. A Simple, Fast, and Accurate Algorithm to Estimate Large Phylogenies by Maximum Likelihood.Rannala B, editor. Syst Biol [Internet]. 2003 Oct 1;52(5):696–704. Available from: http://academic.oup.com/sysbio/article/52/5/696/1681984

32. Simmons MP, Ochoterena H. Gaps as Characters in Sequence-Based Phylogenetic Analyses. Syst Biol [Internet]. 2000 Jun;49(2):369–81. Available from: http://academic.oup.com/sysbio/article/49/2/369/1687092/Gaps-as-Characters-in-SequenceBased-Phylogenetic

33. Espírito-Santo MD, Cubas P, Lousã M, Pardo C, Costa JC. Ulex parviflorus sensu lato (Genisteae, Leguminosae) en la zona centro de Portugal. An del Jardín Botánico Madrid. 1997;55(1):49–66.

34. Soltis PS, Soltis DE. Multiple Origins of the Allotetraploid Tragopogon mirus (Compositae): rDNA Evidence. Syst Bot [Internet]. 1991 Jul;16(3):407. Available from: https://www.jstor.org/stable/2419333?origin=crossref

35. Soltis PS, Plunkett GM, Novak SJ, Soltis DE. Genetic Variation in Tragopogon Species: Additional Origins of the Allotetraploids T. mirus and T. miscellus (Compositae). Am J Bot [Internet]. 1995 Oct;82(10):1329. Available from: http://doi.wiley.com/10.1002/j.1537-2197.1995.tb12666.x

36. Liu B, Davis TM. Conservation and loss of ribosomal RNA gene sites in diploid and polyploid Fragaria (Rosaceae). BMC Plant Biol [Internet]. 2011;11(1):157. Available from: http://bmcplantbiol.biomedcentral.com/articles/10.1186/1471-2229-11-157

37. Rosato M, Moreno-Saiz JC, Galián JA, Rosselló JA. Evolutionary site-number changes of ribosomal DNA loci during speciation: complex scenarios of ancestral and more recent polyploid events. AoB Plants [Internet]. 2015;7:plv135. Available from: https://academic.oup.com/aobpla/article-lookup/doi/10.1093/aobpla/plv135

38. Kovarik A, Pires JC, Leitch AR, Lim KY, Sherwood AM, Matyasek R, et al. Rapid Concerted Evolution of Nuclear Ribosomal DNA in Two Tragopogon Allopolyploids of Recent and Recurrent Origin. Genetics [Internet]. 2005 Feb;169(2):931–44. Available from: http://www.genetics.org/lookup/doi/10.1534/genetics.104.032839

39. Guggisberg A, Bretagnolle F, Mansion G. Allopolyploid Origin of the Mediterranean Endemic, Centaurium bianoris (Gentianaceae), Inferred by Molecular Markers. Syst Bot [Internet]. 2006 Apr 1;31(2):368–79. Available from: http://www.ingentaselect.com/rpsv/cgibin/cgi?ini=xref&body=linker&reqdoi=10.1600/036364406777585937

40. Cubas P, Pardo C. Correlations between chromosomal and morphological characters in subspecies of Ulex europaeus L. (Genisteae, Leguminosae) from the north-west of the Iberian Peninsula. Bot J Linn Soc [Internet]. 1997 Nov;125(3):229–43. Available from: https://academic.oup.com/botlinnean/article-lookup/doi/10.1111/j.1095-8339.1997.tb02256.x

41. Bergsten J. A review of long-branch attraction. Cladistics [Internet]. 2005 Apr;21(2):163–93. Available from: http://doi.wiley.com/10.1111/j.1096-0031.2005.00059.x

42. Borowiec ML, Lee EK, Chiu JC, Plachetzki DC. Extracting phylogenetic signal and accounting for bias in whole-genome data sets supports the Ctenophora as sister to remaining Metazoa. BMC Genomics [Internet]. 2015 Dec 23;16(1):987. Available from: https://bmcgenomics.biomedcentral.com/articles/10.1186/s12864-015-2146-4

43. Paun O, Forest F, Fay MF, Chase MW. Hybrid speciation in angiosperms: parental divergence drives ploidy. New Phytol [Internet]. 2009 Apr;182(2):507–18. Available from: http://doi.wiley.com/10.1111/j.1469-8137.2009.02767.x

44. Rothmaler W. Monographie der Gattung Petrocoptis A. Bre. Bot Jahrb Syst. 1941;72:117–30.

